# Rapid protein-protein interaction network creation from multiple sequence alignments with Deep Learning

**DOI:** 10.1101/2023.04.15.536993

**Authors:** Patrick Bryant, Frank Noé

## Abstract

AlphaFold2 (AF) can evaluate protein-protein interactions (PPIs) with high accuracy by finding evolutionary signals between proteins but comes with a high computational cost. Here, we speed up the prediction with AF for PPI network prediction 40x and reduce the disk space requirements 4000x for a set of 1000 proteins. Our protocol is easy to install and freely available from: https://github.com/patrickbryant1/SpeedPPI.

## Main

Protein-protein interactions (PPIs) govern many cellular processes [1,2] and are essential for understanding and identifying disease targets [3]. Constructing PPI networks is a difficult task due to noisy experimental data. Recently, it has been shown that adaptations of AlphaFold2 [4] (AF) for protein complexes (FoldDock [5]) can rival high-throughput yeast-two-hybrid and mass spectrometry screens in identifying PPIs [6].

Due to the computational requirement of multiple sequence alignment (MSA) creation and structure prediction, it is not feasible to apply high-accuracy methods on a proteome scale. At the same time, single-sequence methods (i.e. those not dependent on an MSA) for PPI prediction have been found to be unreliable [7]. In contrast to other methods, AF for single chains has not been trained on PPIs and, therefore, successful PPI prediction indicates that the network generalizes to this new task. AF has been evaluated for PPI annotation on 1481 known interacting and 5694 non-interacting protein pairs [5]. The score pDockQ (equation 1) was shown to select true PPIs with an area under the ROC curve of 0.87 and only positive PPIs were observed at pDockQ scores above 0.5 [5].

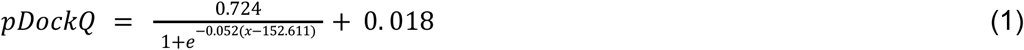

Here, x is the average interface predicted lDDT from the AF predictions multiplied with the logarithm of the number of predicted inter-chain contacts (Cβs within 8Å).

To reduce the computational time and disk space required to annotate an entire PPI network we apply several strategies (Figure 1). **1)** We apply a reduced MSA creation procedure by searching only Uniclust30 compared to the default one of AF [4,5] which searches BFD[8,9][9–11][8,9], MGnify[12] and UniRef90[13]. A slightly increased success rate (fraction of interactions with DockQ>0.23) in the structure prediction of PPIs has been observed using this reduced MSA generation [5]. **2)** We pair MSAs using species information within the prediction run and keep all single-chain information by block diagonalization. **3)** We rewrite the prediction script to prefetch features on the CPU while predictions are running on the GPU, meaning that the total prediction time is that of the GPU processing. Overall, our procedure speeds up PPI network prediction approximately 40 times compared to the default AF procedure for 1000 proteins (499’500 pairwise interactions). The pipeline is compatible with any Unix-based system and can be easily parallelized using the provided source code.

**Figure 1.**
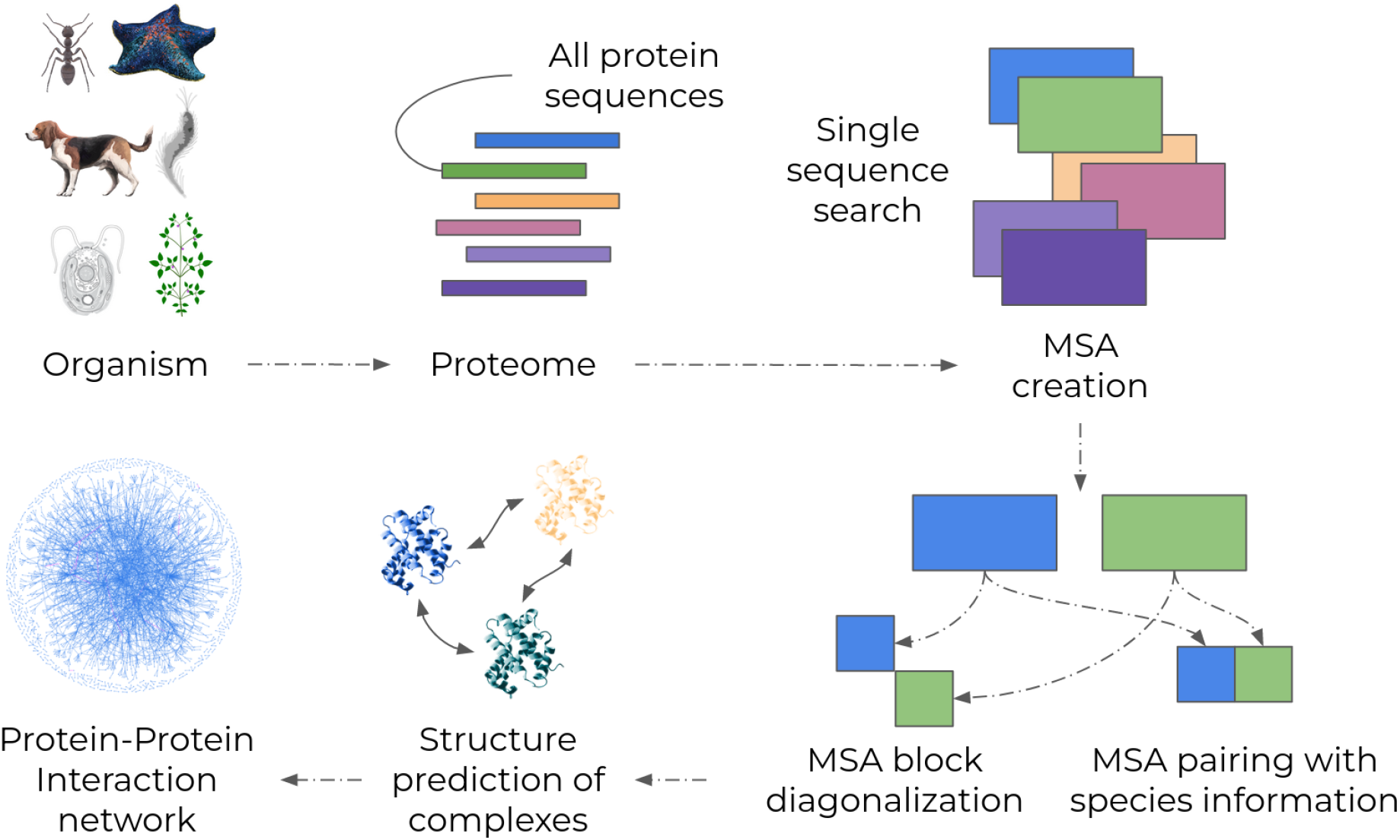
Overview of the procedure used to predict protein-protein interaction (PPI) networks. Given an organism, the proteome is extracted (e.g. from UniProt). All sequences are used in single-sequence mode to create MSAs with HHblits searching only Uniclust30. The MSAs are paired based on species information and all single-chain information is maintained by block diagonalization. The MSA pairing and block diagonalization is done within the network, avoiding writing this information to disk which reduces disk space requirements and the total prediction time. The structure prediction is made with AlphaFold2 and the features are prefetched on the CPU, meaning that the total prediction time is only that on the GPU. The predictions are evaluated with the pDockQ score within the prediction runs and only models above a certain threshold (default = 0.5) are saved to disk. The resulting high-scoring predictions are used to build a PPI network.

We analyse the speedup for different reduction steps calculated for all pairwise interactions from 1000 unique proteins (Figure 2a). The main computational bottleneck for structure prediction is the MSA generation. The reduced MSA creation has a roughly 8-fold speedup compared to the default generation per chain. As all pairs have to be searched in unison using the default AF option with concatenated chains (or AlphaFold-multimer [14]), the speedup of creating MSAs for every single chain and then combining these have a much larger total impact on the computational time (11’395-fold reduction in MSA search time, see Methods). Instead of saving PPI MSAs or features to disk, they are processed within the prediction runs, resulting in a 4’460 -fold reduction in the disk space requirements for a set of 1000 proteins (Figure 2b). The user can set a pDockQ score threshold to determine which structures are saved to avoid saving low-quality predictions.

**Figure 2.**
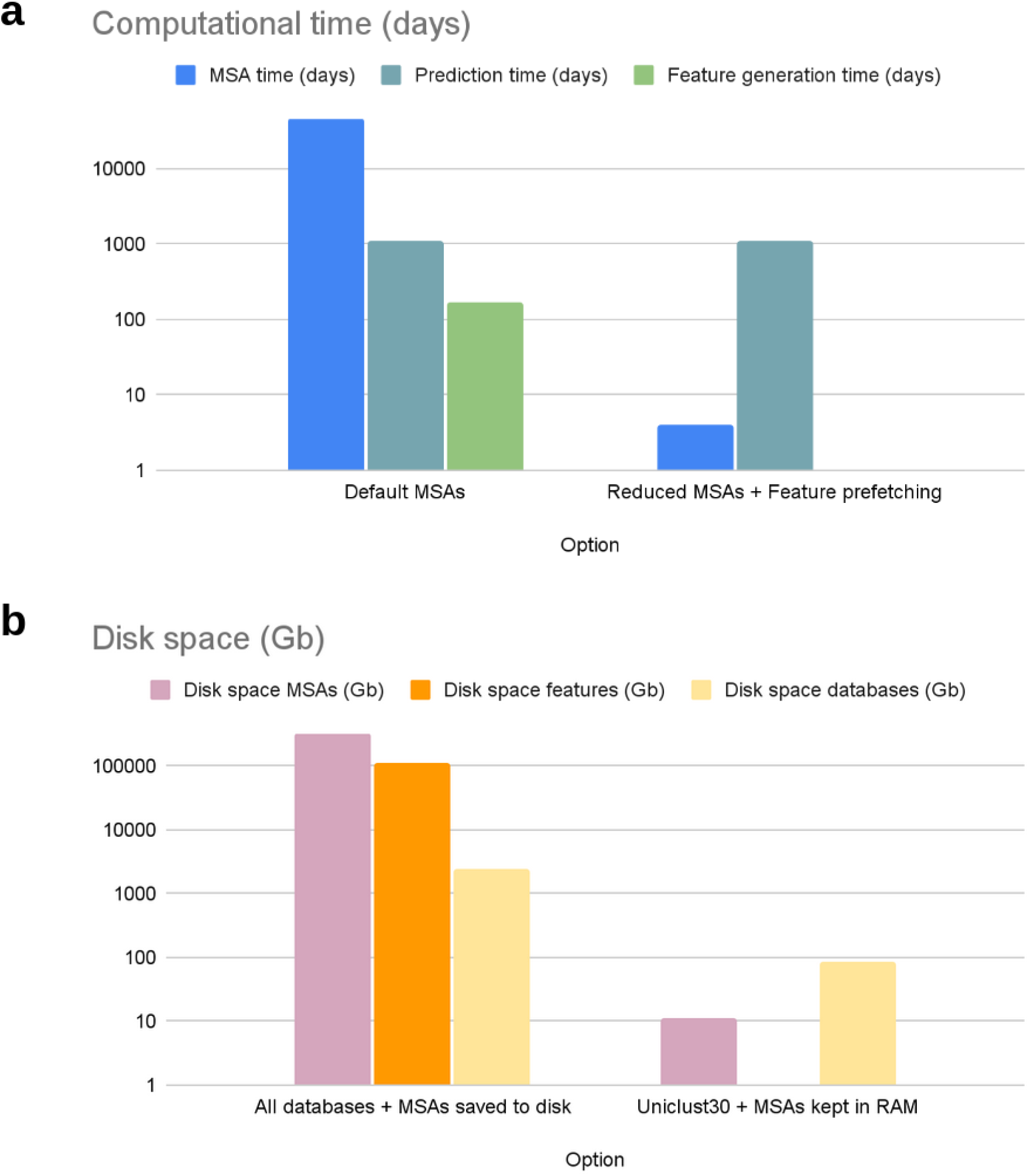
**a)** Computational time (log scale) for different reduction steps calculated for all pairwise interactions over a set of 1000 proteins (499’500 interactions in total). The prediction time here is calculated using an NVIDIA A100 GPU with 40 GB RAM and MSA generation with 16 CPU cores from an Intel Xeon E5-2690v4. The time per unit is taken over a few examples and is displayed as an example. The relative numbers will be similar between the options regardless of the time per example used (which depends on the computational infrastructure). The time requirement is reduced by approximately 40x with the reduced MSA option. The exact number of days is available in Table 1. **b)** Disk space requirements (log scale) for the databases and MSAs. The average disk space per Uniclust30 MSA is 11 Mb and using all databases from AF 650 Mb. Not saving the MSAs or features to disk has the largest effect. In total, the space requirement is reduced by approximately 4000x using Uniclust30 and keeping the MSAs and features in RAM for the predictions only. The exact number of GBs is available in Table 2 (Methods).

The databases used in AlphaFold (BFD[8,9], MGnify[12], PDB70, PDB mmCIF and PDB SEQRES[15], UniRef30 and UniRef90[13]) take a total of 2.49 TB and require hours to days for downloading and processing. Using only Uniclust30 reduces the memory requirement to 87 Gb, a 97% reduction in disk space. This database can be fetched in minutes, making the installation requirements more suitable for a larger user group. The overall memory requirement for the installation of the protocol described here is approximately 90 Gb. Due to the reduction in generated MSA information searching only Uniclust30 with HHblits, the average disk space needed is 11 Mb (1.7%) per MSA compared to 650 Mb for the MSAs generated with AF. No disk space is needed for the features as these are kept in RAM, which takes an average of 220 Mb per complex.

**Table 1.**
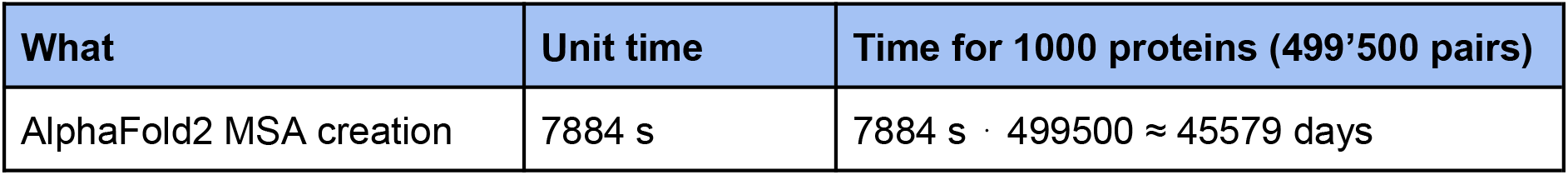
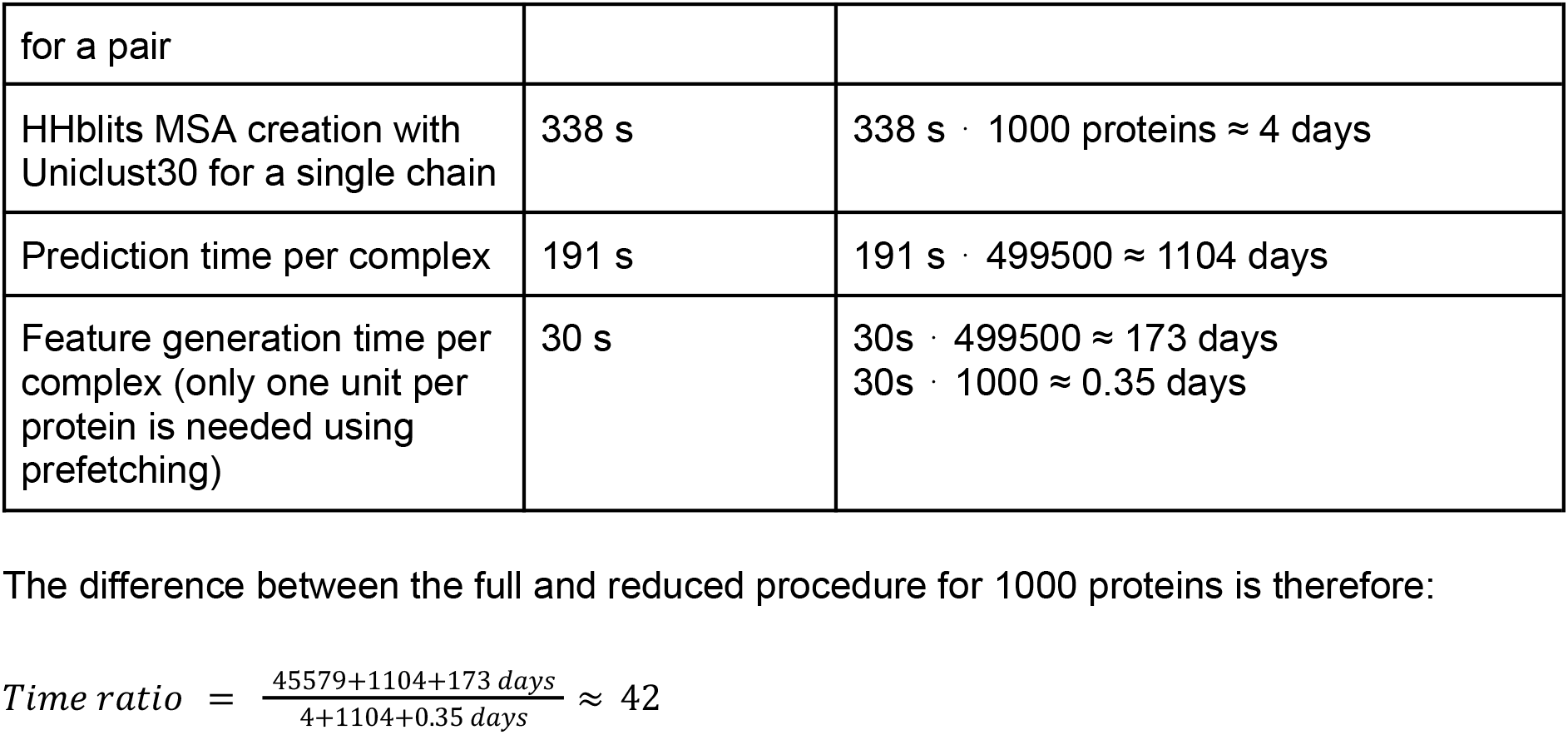
Time for MSA creation and complex prediction for different options.

**Table 2.** Disk space requirements for databases and MSAs using different options.

| What | Unit disk space | Disk space for 1000 proteins (499'500 pairs) |
| --- | --- | --- |
| Disk space for all AlphaFold2 databases | 2490 Gb | 2490 Gb |
| Disk space for Uniclust30 | 87 Gb | 87 Gb |
| MSA disk space per complex using all AlphaFold2 databases | 650 Mb | 324675 Gb |
| MSA disk space per single chain using Uniclust30 | 11 Mb | 11 Gb |
| Feature disk space per interaction | 220 Mb | 109890 Gb |

The difference between the full and reduced procedure for 1000 proteins is therefore:

We provide two options for running the pipeline to suit a variety of users, either all-vs-all or some-vs-some. In the all-vs-all mode, a full PPI network is evaluated for a list of protein sequences (as in a real proteome evaluation setting). In some-vs-some, two lists of proteins of interest can be provided. All proteins in list one will be run against all proteins in list two and a network will be created using all possible pairs between the lists. Note that this evaluation will benefit from a similar speedup since if two lists of 1000 proteins each are analysed there are 1000^2^=1’000’000 possible pairs. We hope that the speedup, reduced disk space and simplifications made here will result in an acceleration in the study of PPIs and new biological insights.

## Methods

### MSA creation and analysis

The most expensive step of the protein structure prediction with AlphaFold2 is the creation of MSAs both in memory consumption and computational time. To speed this process up, we search only Uniclust30[16] with HHblits from HHsuite 3.3.0 [17]) for every single chain. The chains are thereafter paired using the organism identifiers [8] from the hits. This process takes on average 2 seconds, making it negligible compared to the search (average 338 s) with 16 CPU cores from an Intel Xeon E5-2690v4. We note that these timings can be highly variable, but that the relationships observed between the full and reduced searches should be similar.

### Structure prediction and scoring

To use GPU resources more efficiently, we prefetch data on the CPU that are loaded onto the GPU for prediction. Running AlphaFold2 on the same MSAs in the default way took approximately 3 minutes (191 s) per example and with the prefetching approximately 30 s to a few minutes will be saved per example using NVIDIA A100 GPUs with 40 GB RAM. Note that the time saved by the prefetching can be highly variable depending on the bandwidth. With prefetching, features are always ready to go to the GPU, making the prediction process as efficient as possible. The features are kept within the prediction script and never saved to disk, saving on average 220 Mb per example. The same with the scoring of the models using pDockQ. This means that users can set a threshold and only save predictions with a score above a certain threshold, reducing the disk space requirements further and saving a negligible 0.6 Mb per model below the score threshold (not included in the disk space calculations).

### Time complexity and disk space

The number of possible pairwise interactions from a set of n proteins is:

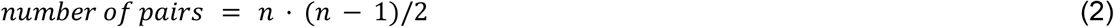

For a set of 1000 proteins (the order of the smallest known proteomes[18]), this results in

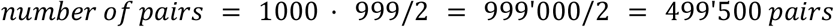

## Data availability

The dataset and metrics used for the creation of FoldDock and pDockQ are available in the original publication [5] and from the GitHub repository which contains this pipeline (https://gitlab.com/ElofssonLab/FoldDock and https://github.com/patrickbryant1/SpeedPPI).

## Acknowledgements

The organism depictions in Figure 1 were created with https://bioicons.com/ and are available under the CC BY 3.0 license (https://creativecommons.org/licenses/by/3.0/).

